# Single-cell Stereo-seq enables cell type-specific spatial transcriptome characterization in *Arabidopsis* leaves

**DOI:** 10.1101/2021.10.20.465066

**Authors:** Keke Xia, Hai-Xi Sun, Jie Li, Jiming Li, Yu Zhao, Ruiying Chen, Guangyu Liu, Zhiyong Chen, Ruilian Yin, Shijie Hao, Jing Wang, Qing Xie, Jiangshan Xu, Yuxiang Li, Ao Chen, Longqi Liu, Ye Yin, Huanming Yang, Jian Wang, Ying Gu, Xun Xu

**Author notes:** Correspondence (X.X.), (Y.G.), (J.W.). These authors contributed equally to this work (Keke Xia, Hai-Xi Sun, Jie Li, Jiming Li, Yu Zhao).

## Abstract

Understanding the complex functions of plant leaves requires spatially resolved gene expression profiling with single-cell resolution. However, although *in situ* gene expression profiling technologies have been developed, this goal has not yet been achieved. Here, we present the first *in situ* single-cell transcriptome profiling in plant, scStereo-seq (single-cell SpaTial Enhanced REsolution Omics-sequencing), which enabled the *bona fide* single-cell spatial transcriptome of *Arabidopsis* leaves. We successfully characterized subtle but significant transcriptomic differences between upper and lower epidermal cells. Furthermore, with high-resolution location information, we discovered the cell type-specific spatial gene expression gradients from main vein to leaf edge. By reconstructing those spatial gradients, we show for the first time the distinct spatial developmental trajectories of vascular cells and guard cells. Our findings show the importance of incorporating spatial information for answering complex biological questions in plant, and scStereo-seq offers a powerful single cell spatially resolved transcriptomic strategy for plant biology.

## INTRODUCTION

Leaves are one of the most important organs for plants. In fully grown leaves, the main leaf cell types include epidermis, mesophyll and vasculature. In *Arabidopsis*, the epidermis is composed of guard cells and trichomes that are embedded in the single cell layer of pavement cells. The upper and lower epidermal cells enclose the palisade and spongy mesophyll, which represent the main photosynthetic capacity of the leaf. Embedded in the mesophyll cells are the vascular cells(Svozil et al., 2015; Tsukaya, 2013). Since individual cell types have specialized functions during leaf development and response to environmental stimuli, a comprehensive *in situ* cell type-specific multi-omics profiling is needed for understanding the complexity of plant leaves. However, this has not yet been achieved.

Recently, high-throughput single cell RNA sequencing (scRNA-seq) technology provides new insights into leaf cell biology, which has facilitated our understanding of stomata and vasculature development at single-cell level(Kim et al., 2021; Liu et al., 2020b). Despite its advantage in single-cell resolution, its application to plant leaf studies still faces challenges(Gurazada et al., 2021). First, protoplasting is still difficult for many plants leaf and other organs, and it also causes changes in the expression of hundreds of genes, which may influence the following transcriptome profiling(Tian et al., 2020). Second, many cell types are resistant to protoplasting, and protoplasting may yield bias proportions of cell types(Bezrutczyk et al., 2021). And third, the most important, the tissue dissociation leads to the loss of spatial information of cells. Spatial information is not only important for the analysing of cell-cell interaction and cell-environment interaction but also crucial for cell type identification, especially for cell types with limited knowledge on marker genes. With spatial information, it could be possible to distinguish cell sub-types and analyze subtle transcriptomic differences between cell sub-types, such as between upper and lower epidermal cells and between palisade and spongy mesophyll cells(Kim et al., 2021; Liu et al., 2020b). Since these drawbacks of scRNA-seq for plant studies, the development of *in situ* gene expression profiling technologies are motivated.

A number of remarkable high-throughput transcriptomic profiling of *in situ* gene expression methodologies have been developed recently(Chen A. et al., 2021; Chen et al., 2015; Cho et al., 2021; Eng et al., 2019; Jiao et al., 2009; Lee et al., 2014; Liu et al., 2020a; Nichterwitz et al., 2016; Rodriques et al., 2019; Srivatsan et al., 2021; Stahl et al., 2016; Stickels et al., 2021; Vickovic et al., 2019; Wang et al., 2018), such as Slide-seq, DBiT-seq (deterministic barcoding in tissue for spatial omics sequencing). Up to now, most of these methods only demonstrate their applications in mammalian systems, and only one method has been applied in plant systems so far (Giacomello et al., 2017). However, the achieved resolution of this approache is still at tissue/domain level, rather than single-cell level. Among all *in situ* gene expression methodologies, Stereo-seq (SpaTial Enhanced REsolution Omics-sequencing) is so far the only sequencing-based spatially resolved transcriptomic technology with subcellular resolution(Chen A. et al., 2021). This technique exhibits a great potential for achieving single cell resolution *in situ* gene expression profiling in plant biology.

To enable *in situ* single-cell level transcriptome analysis in plant leaves, we set out to apply Stereo-seq in *Arabidopsis*. In this study, combining plant cell wall staining with high-resolution spatial transcriptomics using Stereo-seq, we present the first *in situ* single-cell transcriptome profiling in *Arabidopsis* leaves. Single-cell Stereo-seq (scStereo-seq) not only enables us to identify the main leaf cell types, but also allows us to distinguish cell sub-types, including upper and lower epidermal cell, palisade mesophyll cell and spongy mesophyll cell. Furthermore, we also show the existence of cell type-specific spatial gene expression gradients from main vein to leaf edge. We reconstructed those gradients to show for the first time the developmental trajectories of specific cell types according to their spatial distribution.

## RESULTS

### Establishing single-cell Stereo-seq method in *Arabidopsis* leaves

To obtain *in situ* transcriptome profiling at single-cell level, we set out to apply Stereo-seq in *Arabidopsis* leaves. We employed eleven *Arabidopsis* cauline leaf cross sections to explore the applicability of Stereo-seq in the plant system (Figure 1A). *A. thaliana* cauline leaves were cryosectioned and positioned on top of the chip surface with DNA nanoball (DNB) docked in a grid-patterned array of spots, each spot with a size of 220 nm in diameter and a center-to-center distance of 715 nm. The DNB contains random barcoded sequences, the coordinate identity (CID), molecular identifiers (MID) and polyT sequence-containing oligonucleotides, and the polyT was designed to capture mRNAs. After cell wall staining and imaging, the same section was sequenced with Stereo-seq. Finally, eleven leaves with relatively good morphology from two chips were selected for further analysis. To assess whether the detected transcript signal was stringently confined in leaf areas in the bright field, we assessed the molecular identifiers (MID) distribution. Clearly, the MID distribution was specially localized in the leaf areas, and no transcript signal diffusion was observed (Figure 1B, S1A and S1B).

**Figure 1.**
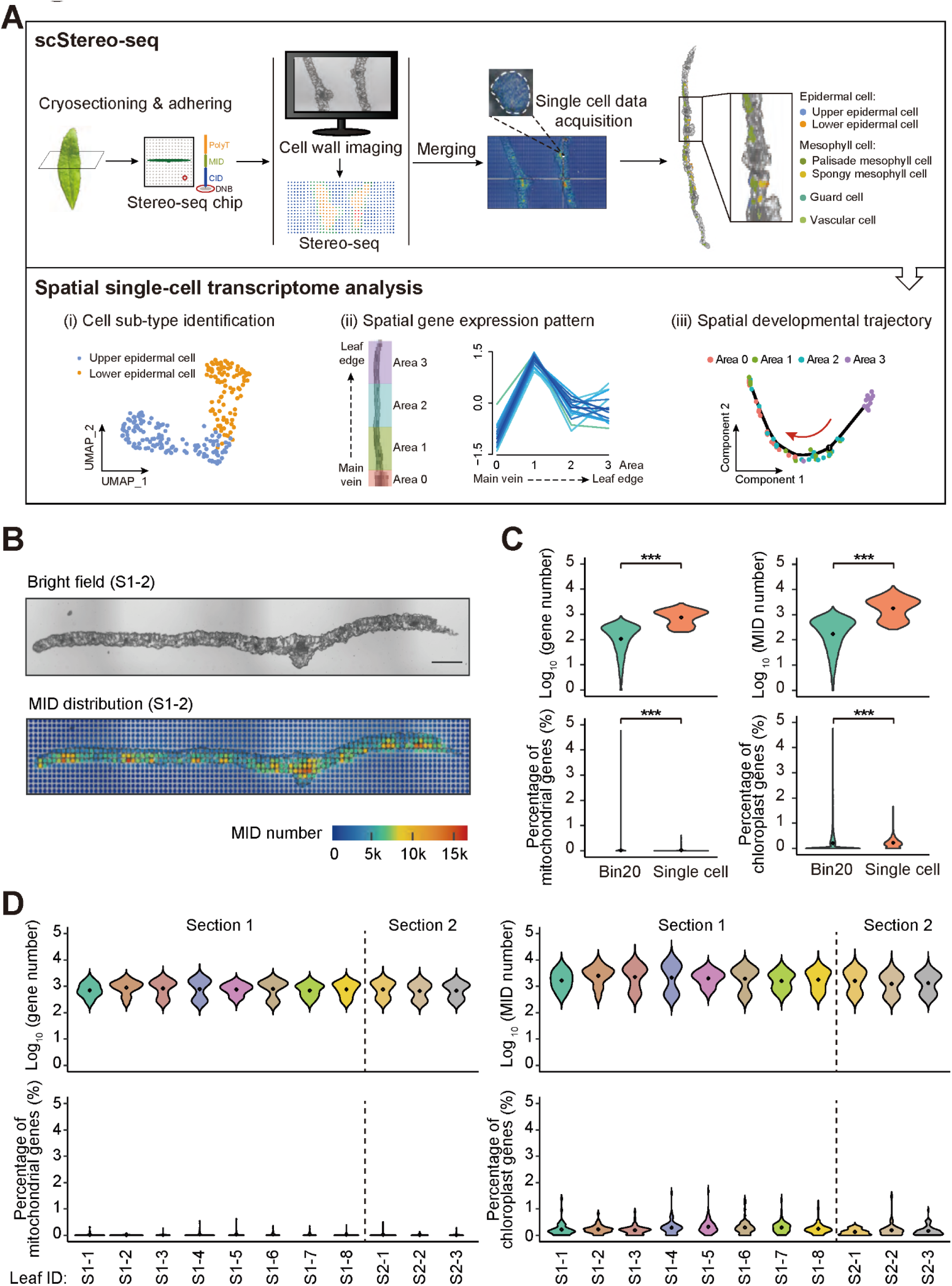
Single-cell level of spatially resolved transcriptome in *Arabidopsis thaliana* cauline leaves. (A) Schematic representation of the single-cell Stereo-seq procedure. *A. thaliana* cauline leaves are cryosectioned and positioned on top of the chip surface with DNA nanoball (DNB) docked in a grid-patterned array of spots, and capture probes contain CID (coordinate identity) barcode, MID (molecular identifiers) and PloyT oligos to enable the recordation of the spatial coordinates, the identification of unique transcripts per gene, and the capture of mRNAs. After cell wall staining and imaging, the same section is sequenced with Stereo-seq. Through the combined high-resolution image and molecular identifiers (MID), single-cell level of MID distribution is achieved. The lasso tool is used to manually extract single cells, and major cell types of cauline leaves are identified and mapped back to the bright field image. Using the spatial single-cell data, several cell sub-types are distinguished (i). And next, the leaf is divided into four distinct parts and spatial gene expression pattern (ii) and spatial developmental trajectory (iii) are determined. (B) A bright field of *Arabidopsis* cauline leaf (leaf #2 of section #1, S1-2) and MID distribution in the leaf. The color bar represents the number of MIDs. The scale bar is 500 μm. (C-D) Violin plots represent the number of genes and MIDs per cell/bin, and the percentage of mitochondria and chloroplast genes per cell/bin of sequenced cells/bins in all cauline leaves (C) and in different cauline leaves (D). The black diamond indicates the median number of genes/MIDs or percentage of mitochondria genes/chloroplast genes per cell/bin. Asterisks indicate statistically significant differences: (***) P < 0.001.

To visualize cell-cell boundary, we performed toluidine blue (TB) staining of plant cell wall on the same section we used for Stereo-seq. After merging the staining image and sequencing data in the Stereomics visualization system, we then used lasso tool to manually extract single-cells based on cell wall boundary and coordinated them with MID counts to obtain specific data for each single cell. For eleven leaf samples, we obtained 1,177 cells in total, and these cells had clear cell boundaries and could be clearly assigned to specific cell types. After filtering (gene number: 200-5000; percentage of mitochondrial gene: < 10%; see Methods), a total of 871 high-quality single cells were subject to further analyses. We then determined the gene number, the MID number, the percentage of mitochondria genes, and the percentage of chloroplast genes in each cell to evaluate our data quality. The majority cells had similar gene numbers and MID numbers with a strong positive correlation (Figure 1C and S1C), and there was no obvious difference among the eleven leaves (Figure 1D), providing clear evidence of consistency and reproducibility. In addition, when we compared the data obtained by single-cell method mentioned above with that obtained by the standard Stereo-seq data analysis method (the binning method using 20 x 20 DNBs, hereafter referred to as ‘Bin20’) (Chen A. et al., 2021), we observed a significant increase in gene and MID numbers of the single-cell method, indicating that single cell extraction greatly improved the data quality of Stereo-seq (Figure 1C). Taken together, we obtained spatially resolved single-cell transcriptome profiling of *Arabidopsis* cauline leaves.

### Obtaining cell type-specific transcriptomes at *bona fide* single-cell level

With morphology features and their locations in the leaves, the 871 cells were classified into 226 epidermal cells (130 upper and 96 lower epidermal cells), 472 mesophyll cells (331 palisade and 141 spongy mesophyll cells), 95 guard cells and 78 vascular cells (Figure 2A and Table S1). The gene number, UMI number, percentage of mitochondria genes and the percentage of chloroplast genes per cell showed similar patterns in four different major cell types (Figure 2B), demonstrating that our approach was able to obtain unbiased single-cell transcriptome of all major cell types of *Arabidopsis* leaves.

**Figure 2.**
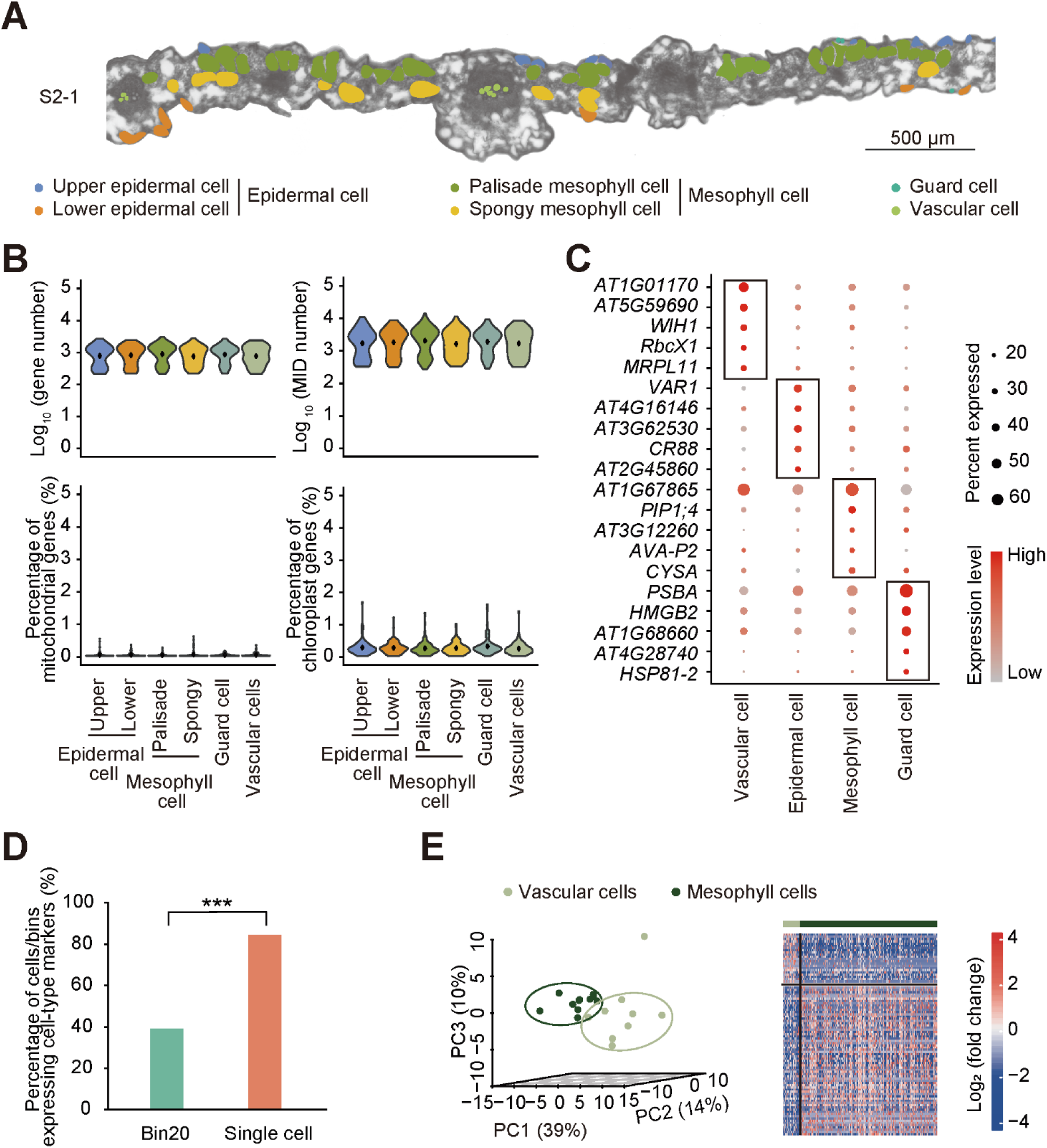
Technical reproducibility and transcriptome diversity of cell types from different leaves. (A) Distribution of different cell types in the bright field picture. The scale bar is 500 μm. (B) Violin plots show the gene number, the MID number, the percentage of mitochondria genes, and the percentage of chloroplast genes per cell in different cell types. The black diamond indicates the median number of genes/MIDs or percentage of mitochondria genes/chloroplast genes per cell. (C) Dot plots showing the top five up-regulated marker genes of each cell type. The circle size indicates the percentage of cells expressing the marker genes, while color represents expression value. (D) Percentage of cells/bins expressing cell-type markers. (E) PCA plot of vascular cells and mesophyll cells in different leaves (left) and heatmap showing DEGs between vascular cells and mesophyll cells (right). Blue and red represent log2-transformed fold change < 0 and > 0, respectively. Asterisks indicate statistically significant differences: (***) P < 0.001.

In order to confirm the reliability of cell type-specific transcriptome, we performed the marker genes, Principle Component Analysis (PCA), and differentially expressed genes (DEGs) analyses among the four major cell types. All four major cell types expressed cell-type-specific marker genes (top five up-regulated genes were listed) (Figure 2C). Meanwhile, the percentage of cells expressing cell-type markers of single-cell method was significantly higher than those of Bin20 (Figure 2D and S2A-D), showing that the single-cell method could better reflect the cell-type-specific transcriptome characteristics. As a result, PCA and differential expression analyses using the single-cell transcriptome revealed that the same cell type from different leaves was clustered together while different cell types from different leaves were separately grouped (Figure 2E and S2E-I). These significant differences among the four major cell types demonstrated the consistency and reproducibility of scStereo-seq data at cell-type level.

We further validated the reliability of the four cell types we distinguished in two aspects. On the one hand, we detected the expression level of the previously reported marker genes in each cell type, *GSTU20*, *AT2G45860*, *AT5G53940*, and *HSP81-2,* which were specifically expressed in vascular cells, epidermal cells, mesophyll cells and guard cells, respectively(Kim et al., 2021; Liu et al., 2020b; Zhang et al., 2021). And the results showed that these genes were highly expressed in their corresponding cell types and lowly expressed in other cell types (Figure 3A and S3A). In addition, these marker genes were specifically expressed in their corresponding loci in the TB staining images (Figure 3A and S3A). On the other hand, we investigated the biological processes enriched in each cell type, and found that all cell types were enriched in biological processes corresponding to their specialized functions (Figure 3B). For instance, vascular cells were enriched in transportation-related pathways^38,39,^ while mesophyll cells were enriched in photosynthesis-related pathways and sucrose metabolic process(Ruan, 2014). Three photosynthesis-related Gene Ontology (GO) terms were also found in vascular cells, and this feature has been experimentally validated in the vascular tissues of phylogenetically widespread C3 plants(Brown et al., 2010; Gao et al., 2018; Hibberd and Quick, 2002). These results demonstrated the reliability of cell type identification using extracted single-cell data using scStereo-seq. So far, we obtained *in situ* reliable cell type-specific transcriptome data at single-cell level, for the first time.

**Figure 3.**
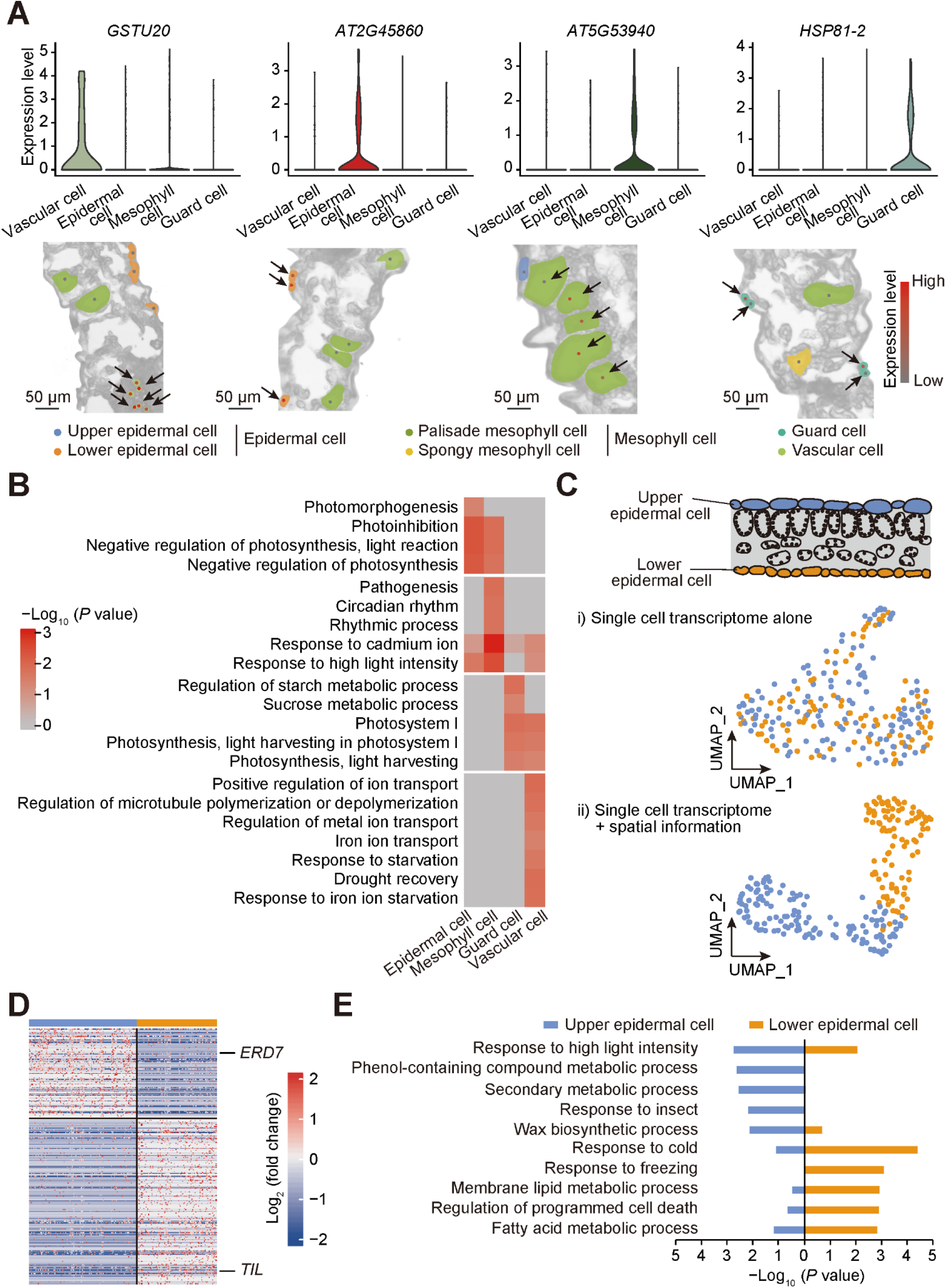
Validation and investigation of transcriptome characteristics of cell types and cell sub-types. (A) Validation of cell types using cell-type-specific marker genes (top panel). Representative images showing expression levels of marker genes in the corresponding cells in the bright field (lower panel). (B) Enriched GO terms for four different cell types. (C) scStereo-seq allows separation between upper and lower epidermal cells. Distribution of upper and lower epidermal cells: i) without spatial information (using variable genes of all epidermal cells); and ii) with spatial information (using DEGs between upper and lower epidermal cells). Blue dots represent upper epidermal cells and orange dots represent lower epidermal cells. (D) DEGs between upper and lower epidermal cells. Blue and red represent log2- transformed fold change < 0 and > 0, respectively. (E) GO enrichment analysis of DEGs between upper and lower epidermal cells.

### Spatial information and cell boundary are necessary to identify subtle transcriptional differences between cell sub-types

For *Arabidopsis* leaves, epidermis can be subdivided into upper epidermal cell and lower epidermal cell, and mesophyll cell can also be subdivided into spongy mesophyll cell and palisade mesophyll cell. However, because of highly similar transcriptomes between cell sub-types, scRNA-seq has not been able to distinguish these cell sub-types in previous studies(Kim et al., 2021; Liu et al., 2020b). In this study, using scStereo-seq, we could extract single cells and classify cell types based on TB staining of cell walls and spatial histological information. Therefore, we were able to distinguish upper and lower epidermal cells, as well as spongy and palisade mesophyll cells: Without the spatial histological information, cell sub-types could not be distinguished using single cell transcriptome alone (Figure 3C and S3B); without the accurate cell boundaries (the Bin20 method), they could not be distinguished either (Figure S3E and S3F). Because inaccurate cell boundaries conferred poor data quality (Figure 1C) and fewer transcriptomic features (Figure 2D), we demonstrated that both spatial coordinate and accurate cell boundaries were required for cell sub-type classification in plant leaves, and also showed the necessity of using scStereo-seq technology for complex plant tissue studies.

With obtained datasets and DEGs of these cell sub-types, we further investigated their functional differences, which has significant biological implications and has not yet been achieved so far. As shown in Figure 3E, biological processes associated with high light response, wax biosynthetic process and response to insect were enriched in upper epidermal cell, while biological terms related to freezing, membrane lipid metabolic process and regulation of programmed cell death, were significantly enriched in lower epidermal cell (Figure 3E). For instance, *ERD7* is upregulated in various biological processes, including response to high light(Kimura et al., 2003), whereas *temperature-induced lipocalin* (*TIL*) is involved in cold stress response(Frenette Charron et al., 2002; Kawamura and Uemura, 2003; Miki et al., 2019). In this study, using scStereo-seq, we further discovered that *ERD7* and *TIL* were highly expressed in upper epidermal cell and lower epidermal cell, respectively (Figure 3D), indicating upper and lower epidermal cells probably were functionally different in response to high light and cold stress. For mesophyll cell sub-types, we also found that oxygen-related processes were enriched in spongy mesophyll cells (Figure S3D). Taken together, by distinguishing cell sub-types, we revealed the subtle transcriptional differences between upper and lower epidermal cells, spongy and palisade mesophyll cells, which would enable us to discover the complex regulation mechanisms during leaf development and in response to environmental stimuli, in future studies.

### Photosynthesis-related genes exhibited expression gradient from main vein to leaf edge

Genes specifically expressed in certain areas of leaves play key roles in various plant biological processes, such as leaf development and plant photomorphogenesis(Martinez et al., 2021; Tian et al., 2019; Wang et al., 2020). As spatial information of individual cells was documented by scStereo-seq, we were able to assess gene expression levels in different parts of leaves at single cell level, for the first time. As shown in Figure 4A, the main vein area was designated as Area0, and the Area on the one side of the main vein was divided into 3 areas, which were designated as Area 1, 2 and 3, and Areas 1 to 3 gradually away from the main vein. We then determined gene expression patterns in the designated areas of the leaves.

**Figure 4.**
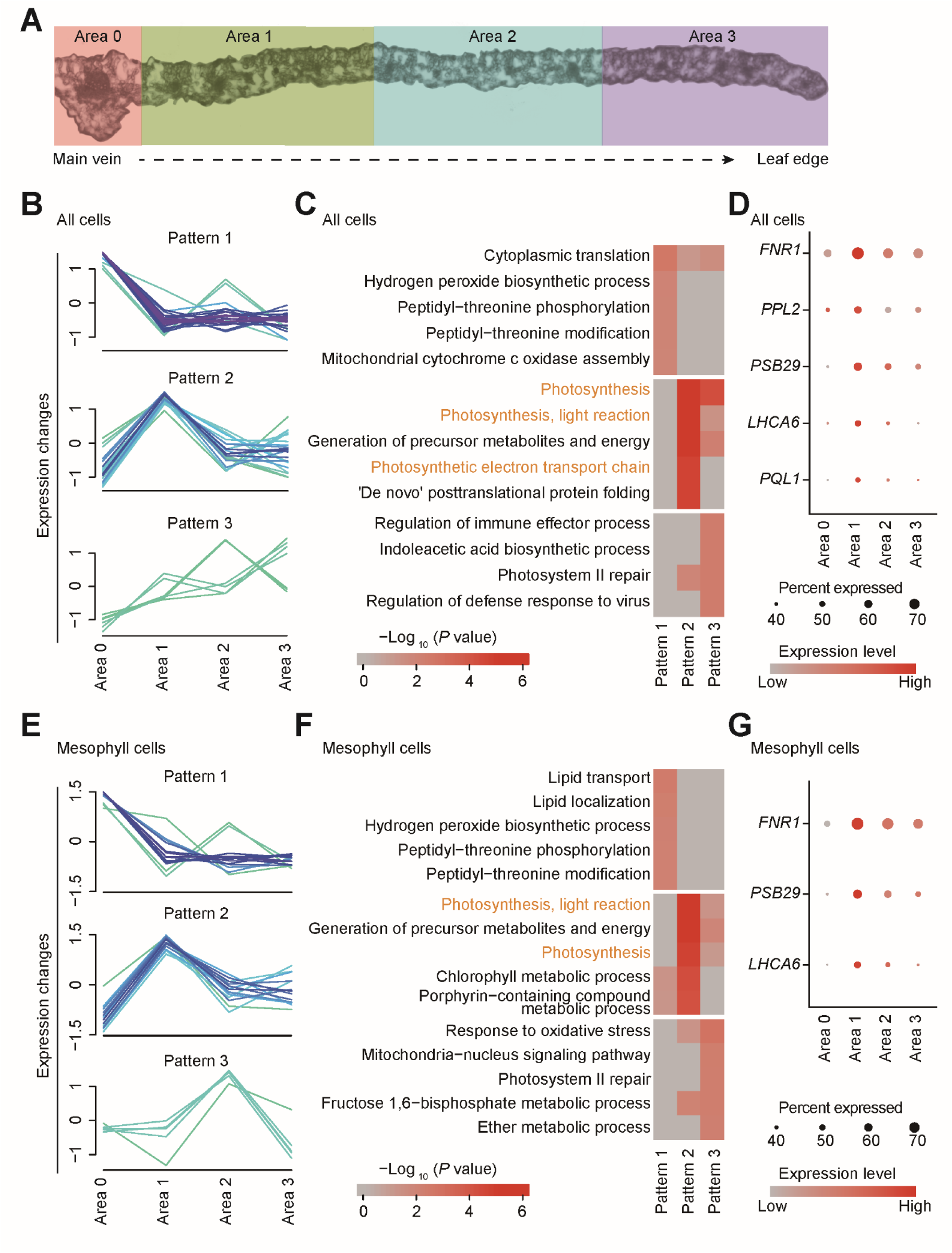
Spatial distribution of photosynthesis-related genes in *Arabidopsis* leaves. (A) Four unique areas (Area 0, 1, 2, 3) from main vein to leaf edge in a leaf. (B) Gene expression patterns for all cells. (C) GO enrichment analysis for genes present in each pattern. (D) Dot plot showing expression levels of representative photosynthesis-related genes in all cells. (E) Gene expression patterns for mesophyll cells. (F) GO enrichment analysis for genes present in each pattern. (G) Dot plot showing expression levels of representative photosynthesis-related genes in mesophyll cells. For dot plots the circle size indicates the percentage of cells expressing the marker genes, while color represents expression value. For pattern analysis, only genes with membership > 0.41 are shown.

Interestingly, 3 patterns of spatial gene expression gradients from main vein to leaf edge were observed when we performed gene expression pattern analysis along the leaf for all cells together (Figure 4B). Pattern 1 genes were highly expressed in the main vein (Area 0) and had a low, relatively stable expression in Area 1, 2 and 3. Pattern 2 genes were highly expressed in Area 1, and the expression level of Pattern 3 genes increased gradually from Area 0 to 3. We found that photosynthesis-related biological processes, such as photosynthesis, photosynthesis electron transport chain and photosynthesis (light reaction), were highly enriched in Pattern 2 (Figure 4C), suggesting that the photosynthetic efficiency decreases gradually from the middle to the edge of the leaf. Given that the photosynthesis process mainly takes place in mesophyll cells and to further validate this observation, we performed the same analysis on mesophyll cell. Three gene expression patterns similar to those of all cells were observed (Figure 4E), and consistently, the photosynthesis-related biological processes were also highly enriched in Pattern 2 in mesophyll cells (Figure 4F). Representative genes were presented, including *PQL1*, *LHCA6*, *PSB29*, *PPL2* and *FNR1* (Figure 4D and 4G), which are known regulators of photosynthesis process(Ishihara et al., 2007; Keren et al., 2005; Morales et al., 2000; Peng et al., 2009; Yabuta et al., 2010). These results further supported that photosynthesis process exhibited a spatial distribution in plant leaves and the photosynthetic efficiency might decrease gradually from the middle to the edge of the leaf. The elevated expression levels of photosynthesis-related genes in Area 1 indicated that photosynthesis was involved in a sophisticated and complex regulation mechanism in leaves, and future studies are needed to uncover the spatial gene regulatory mechanisms.

### Vascular cells and guard cells exhibit distinct spatially resolved developmental trajectories

It isknown that the differentiation at the margin of a leaf is decelerated relative to the more medial regions(Martinez et al., 2021). However, it is still unknown whether all cell types in leaves follow the same development pattern. Here, to address these questions and taking advantage of spatial information of scStereo-seq, we performed pseudotime analysis on vascular cell and guard cell, and compared their developmental stages in the 4 areas as described in last section (Figure 4A).

Pseudotime analysis on vascular cells showed one major trajectory with vascular cells from different leaf areas separately located along the pseudotime path (Figure 5A). Vascular cells from Area 3, the relatively young area in leaves, were distributed at a relatively early pseudotime stage, while vascular cells from Area 2, 1 and 0 were distributed to the end of pseudotime branch (Figure 5A), suggesting that the vascular cells from Area 3 were newly formed relative to the cells from Area 2, 1, and 0. These results showed that the development of vascular cells at the margin of a leaf is decelerated relative to the more medial regions.

**Figure 5.**
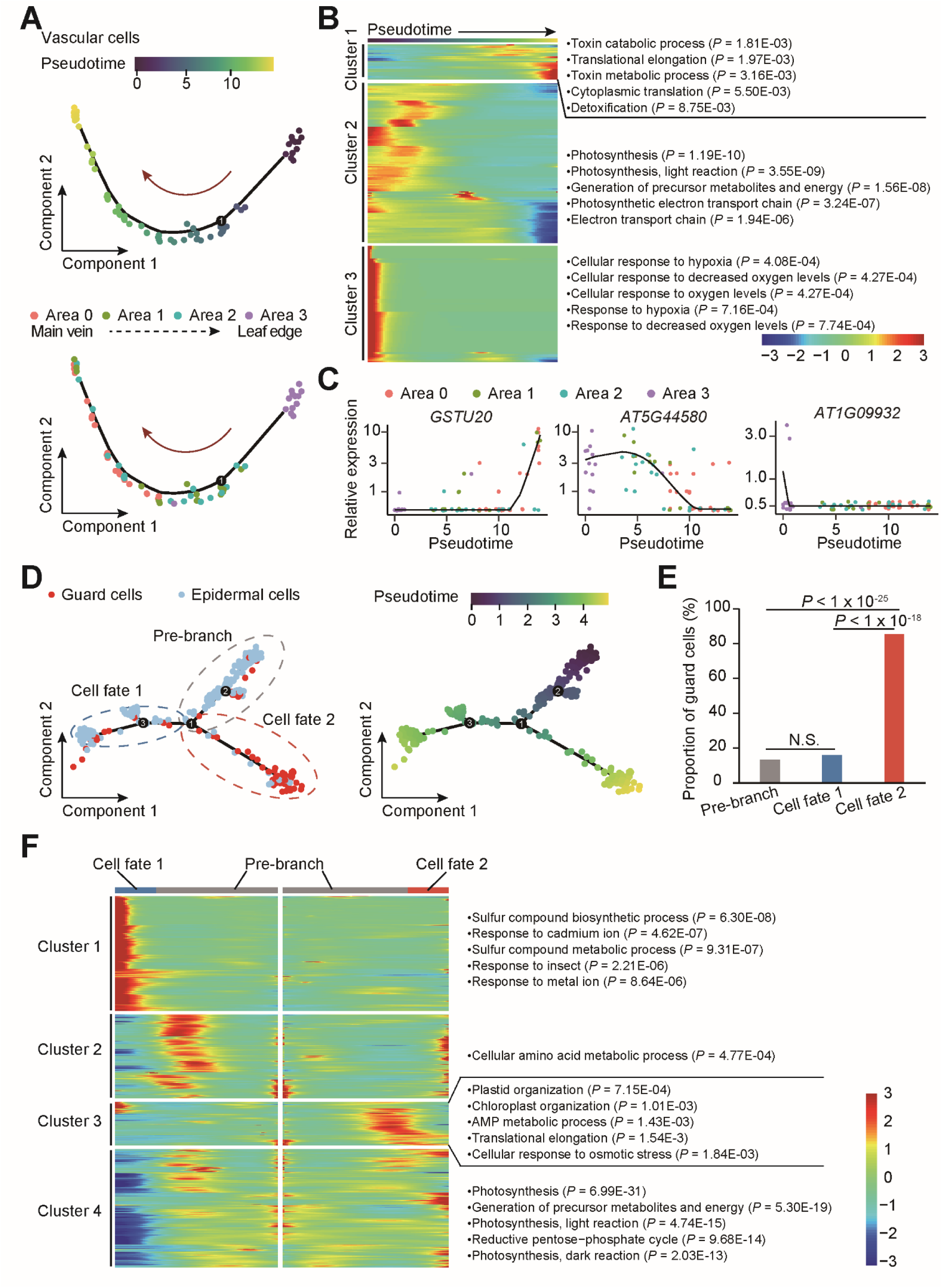
Spatially resolved developmental trajectories of vascular cells and guard cells. (A) Distribution of vascular cells on the pseudotime trajectory (top); distribution of vascular cells from each area on the pseudotime trajectory (bottom). (B) Clustering of differentially expressed genes along a pseudo-time progression of vascular cells (left); top five enriched GO terms for each cluster (right). (C) Gene expression kinetics along pseudotime progression for representative genes; each dot represents one cell and dot color represents its location. (D) Distribution of guard cells and epidermal cells on the pseudotime trajectory branches, including pre-branch, cell fate 1 and cell fate 2. (E) Proportion of guard cells in each branch. (F) Clustering of differentially expressed genes along a pseudotime progression of epidermal cells and guard cells (left); representative enriched GO terms for each cluster (right). N.S. represents not significant.

Moreover, to further investigate the functions of cells in different developmental stages, we assessed gene expression pattern of DEGs along the pseudotime (Figure 5B). DEGs were classified into three distinct clusters. Cluster 1 genes were highly expressed genes at the late stage of the pseudotime, corresponding to cells in the main vein area, and their functions were mainly involved in toxin-related biological processes via GO. Cluster 2 genes that were mainly upregulated in the middle stage of pseudotime were related to photosynthesis, indicating that the photosynthesis process may play significant roles in the development of cells from Area 1 or 2. For cluster 3, responses to oxygen were enriched at the early stage of the pseudotime, corresponding to the leaf edge. Genes involved in vascular development have been studied extensively before(Ischebeck et al., 2013), but their spatial-temporal expression patterns have not been clearly determined. Thus, we determined the pseudotime expression profiles for representative genes from different clusters (Figure 5C). *GSTU20*, *AT5G44580* and *AT1G09932* are probably involved in lignin biosynthesis(Geng et al., 2019). *GSTU20* mainly expressed in Area 0, corresponding to the early formed main vein, while *AT1G09932* mainly expressed in Area 3, corresponding to the latest formed cells, and *AT5G44580* mainly expressed in cells form Area 1 and 2. And several other genes involved in vasculature development also showed spatial-temporal expression changes along the pseudotime (Figure S4A). These results indicated that the maturation of vascular cells went through several development stages, and complex spatial-temporal gene expression regulation was involved in these development stages. In future studies, to better reveal the complex regulatory network of vascular development, it is necessary to conduct independent studies at various development stages.

Because epidermal cell and guard cell differentiate from the same basal epidermal cell(Glover et al., 2016), we investigated the spatial developmental trajectory of epidermal cell and guard cell together. As shown in Figure 5D, epidermal cells and guard cells are located on different branches with one branch mainly containing epidermal cells, designated as cell fate 1, and another branch, designated as cell fate 2, predominantly containing guard cells. More than 80% cells on cell fate 2 were guard cells, which was significantly higher than those on pre-branch and cell fate 1 (Figure 5E). However, when adding spatial information to the development trajectory, we did not find a clear spatial distribution pattern for cells from pre-branch and fate 1 and 2: no obvious difference in spatial distribution of these cells from Area 0 to 3 (Figure S4B and S4C). These data suggest that the development of vascular cells, but not guard cells, at the margin of leaf is decelerated relative to the more medial regions. The development stages with spatial information revealed here could provide important references for future studies on special development stages of vascular cells.

## DISCUSSION

Leaves are the main photosynthetic organs, and they are also involved in responses to biotic and abiotic stresses(Maugarny-Cales and Laufs, 2018). Different cell types in plant leaves exhibit various molecular signatures and perform specific functions. Cell type-specific characterizations are essential for better understanding of the developmental mechanisms of leaves and their responses to environmental stimuli. With the advantages of scStereo-seq that was developed in this study, we successfully revealed the complex cell type-specific and spatial-temporal gene expression features of *Arabidopsis* leaves.

Though plant cell wall is considered to bring difficulties in cell isolation for most plant single-cell studies, our study innovatively uses it for accurate single cell extraction by combining plant cell wall staining with Stereo-seq to obtain well-displayed cell-cell boundaries. The single-cell extraction method displayed a great improvement in data quality when compared with the Standard Stereo-seq Bin20 method. This enables us to generate the first *bona fide* single-cell spatial transcriptome profile in plants and to clearly distinguish cell types and cell sub-types. Although laser capture microdissection(Martinez et al., 2021), MERFISH (multiplexed error-robust FISH)(Xia et al., 2019) and 1cell-DGE (single cell-digital gene expression)(Kubo et al., 2019) can also perform single cell-types and single cell profiling with spatial information in plants, these methods suffer from low throughput and require special equipment which cannot be easily accessed in regular labs. In contrast, transcriptome signals of cells from eleven leaf sections were easily captured on chips in one time using our method, and the high-throughput *in situ* single cell transcriptome profile method allowed us to conveniently capture cells in batch and without the need of special equipment. Although the classification of epidermal cell, mesophyll cell, vascular cell, and guard cell types have been achieved using single cell transcriptomes by scRNA-seq in previous studies(Liu et al., 2020b; Lopez-Anido et al., 2021), identification cell sub-types such as upper and lower epidermal cell, and spongy and palisade mesophyll cell, have not been well achieved so far. In this study, we demonstrate that using single cell transcriptomes alone or the standard Bin20 method could not distinguish these cell sub-types, most likely because these cell sub-types have highly similar transcriptome characteristics and the lacking of cell type-specific marker genes. In contrast, combining spatial information with single cell transcriptomes, we successfully identified these cell sub-types and obtained the cell type-specific characterizations. Thus, scStereo-seq can be easily extended to a variety of model or non-model plant species, avoiding the need of well-established marker genes for highly similar cell types identification.

Furthermore, using scStereo-seq, *bona fide* single-cell transcriptome data could be obtained without preparing protoplasts, thus avoiding the effects on gene expression and biased cell capture(Rich-Griffin et al., 2020; Shaw et al., 2021). The obtained dataset may better reflect the real transcriptional characteristics of the cell types in leaves. For high-throughput single-cell transcriptome analysis in *Arabidopsis* leaves, a large number of single protoplasting cells are needed for capturing rare cell types(Kim et al., 2021; Liu et al., 2020b; Lopez-Anido et al., 2021). And for scRNA-seq, optimized methods might be needed for capturing special cell types(Kim et al., 2021). These problems can be avoided by using scStereo-seq technology.

Based on cell spatial information, we found the existence of cell type-specific spatial gene expression gradients from main vein to leaf edge, such as *FNR1*, *PSB29*, and *LHCA6* in mesophyll cells, which are important components of photosynthesis(Ishihara et al., 2007; Keren et al., 2005; Morales et al., 2000; Peng et al., 2009; Yabuta et al., 2010). The extremely low expression of photosynthesis-related genes in the main vein region (Area 0) is consistent with the main function of leaf vasculature, which plays a key role in solute translocation rather than in photosynthesis, reflecting the reliability of the spatial gene expression pattern. The expression levels of *FNR1*, *PSB29*, and *LHCA6* were extremely high in the leaf area next to the main vein (Area 1), and decreased gradually from the middle to the edge of the leaf, indicating that the area next to the main vein might have a higher photosynthesis capacity compared with leaf edge. In addition, the significance of this spatial expression pattern and the precise spatial regulatory mechanism need to be further investigated. Meanwhile, we reconstructed those gene expression gradients to show for the first time the developmental trajectories of vascular cells and guard cells according to their spatial distribution. The tracking of spatial location of cells at various developmental stages is meaningful. For example, spatial information combined cell type-specific trajectory analysis could provide a reference for assessing the reliability of development trajectory analysis results, and could facilitate researchers to sample cells at specific developmental stages.

The scStereo-seq technology enabled us to reveal the complex cell type-specific and spatial-temporal gene expression features of *Arabidopsis* leaves. However, it is still in its infancy. First, the 871 single cells were manually extracted one by one using lasso tool in this study, which was time- and labor-consuming. Therefore, we are also developing cell boundary algorithms for automatic cell capture to meet the needs of high-throughput cell extraction for large tissues in future studies. Second, improvements in sectioning are required to obtain high-quality cell wall staining images. Because of the high water content in plants, cryosectioning is challenging for many plants or tissues. A high-quality slide is critical to identify all cell types in tissues. In this study, if high-quality slides were available, we were able to identify and extract multiple cell types of veins, which consist at least seven different cell types(Kim et al., 2021). Third, further improvement in the resolution of the stereo-seq chips could facilitate identify tiny cells in certain tissues, such as plant shoot apex.

Taking advantage of the *bona fide* single-cell spatial transcriptome, scStereo-seq has great potential for future studies. For example, based on the high-resolution of scStereo-seq, we would be able to analyze transcriptomes at subcellular level. Due to the simplicity of scStereo-seq compared with MERFISH(Xia et al., 2019) and FISSEQ (fluorescent in situ RNA sequencing)(Lee et al., 2014), it could be applied to broad plant research areas. We can also use scStereo-seq to analyze how different cell types respond to pathogen infection in plant leaf and root, and how response signals are transmitted to distant cells and tissues. Moreover, scStereo-seq can be used to analyze the leaf differences between C3 and C4 plants. By comparing transcriptome differences among bundle sheath cells, other vascular cells and the surrounding mesophyll cells in C3 and C4 plants and constructing developmental trajectories of bundle sheath cells, we could provide insights for transforming C3 plants into C4 plants. In addition, scStereo-seq is also an ideal technology to facilitate building Plant Cell Atlas(Rhee et al., 2019). The scStereo-seq technology developed in this study offers a powerful single cell spatially resolved transcriptomic strategy for systematic studies of plant biology, and it will enable us to understand plants at unprecedented resolution.

## METHODS

### Plant growth condition and tissue collection

*Arabidopsis* (Col-0) seeds were sown and incubated at 25 °C in the photoperiod of 16 hours light / 8 hours dark after surface sterilization with 8% sodium hypochlorite in 0.1% triton X-100 solution. Fresh leaves were collected from five-week-old seedlings and immediately fixed in Carnoy’s fluid (3 ethanol: 1 acetic acid) for 20 minutes. After 10% glycerin treatment, tissues were embedded in cold OCT (Sakura) and stored at -80 °C until processed.

### Stereo-seq chip structure

The Stereo-seq capture chips were used in this study(Chen A. et al., 2021). To generate the DNA nanoball (DNB) array for *in situ* RNA capture, we first synthesized random 25-nt CID (coordinate identity) -containing oligos, circularized with T4 ligase and splint oligos. DNBs were then generated by rolling circle amplification and were loaded onto the patterned chips (65 mm × 65 mm). The chip surface consists of DNA nanoball (DNB) containing random barcoded sequences, the coordinate identity (CID), molecular identifiers (MID) and polyT sequence-containing oligonucleotides. The DNBs are docked in a grid-patterned array of spots, each spot being approximately 220 nm in diameter and with a center-to-center distance of 715 nm. Next, to determine the distinct DNB-CID sequences at each spatial location, single-end sequencing was performed using sequencing primers in a MGI DNBSEQ-Tx sequencer with sequencing strategy SE25. Finally, MID and polyT-containing oligos were hybridized and ligated to the DNB on the chip. This procedure produces capture probes containing a 25 bp CID barcode, a 10 bp MID and a 22 bp polyT ready for *in situ* capture.

CID sequences together with their corresponding coordinates for each DNB were determined using a base calling method according to manufacturer’s instruction of DNBSEQ™ sequencer. After sequencing, the capture chip was split into smaller size chips (10 mm × 10 mm). At this stage, all duplicated CID that corresponded to non-adjacent spots were filtered out.

### Cryosectioning, fixation, staining and imaging

The pre-frozen leaf tissues in OCT were transversely sectioned at 10 µm thickness using a Leika CM1950 cryostat. Tissue sections were adhered to the Stereo-seq chip surface and incubated at 37 °C for 3 minutes. Then, tissues were fixed in methanol and incubated at -20 °C for 30 minutes. The same tissue sections were stained with toluidine blue for stereo-seq, while tissue sections adjacent to those were adhered to glass slides for histological examination using the same staining method. Imaging for both procedures was performed with a Ti-7 Nikon Eclipse microscope.

### Permeabilization, reverse transcription, tissue removal and cDNA release, Library preparation and sequencing

These processes were performed according to the previously reported Stereo-seq method(Chen A. et al., 2021). Tissue patches on the chip were permeabilized using 0.1% pepsin (Sigma, P7000) in 0.01 M HCl buffer (pH = 2), incubated at 37 °C for 10 minutes and then washed with 0.1x SSC buffer (Thermo, AM9770) supplemented with 0.05 U/μL RNase inhibitor (NEB, M0314L) after toluidine blue in tissues was rinsed off with 80% ethanol. Released RNA from permeabilized tissues was captured by the DNB and reverse transcribed overnight at 42 °C using SuperScript II (Invitrogen, 18064-014, 10 U/μL reverse transcriptase, 1 mM dNTPs, 1 M betaine solution PCR reagent, 7.5 mM MgCl2, 5 mM DTT, 2 U/μL RNase inhibitor, 2.5 μM Stereo-TSO and 1x First-Strand buffer).

After *in situ* reverse transcription, tissue patches were washed twice with 0.1x SSC buffer and digested with Tissue Removal buffer (10 mM Tris-HCl, 25 mM EDTA, 100 mM NaCl, 0.5% SDS) at 37 °C for 30 minutes. cDNA-containing chips were then subjected to Exonuclease I (NEB, M0293L) treatment for 3 hours at 37 °C and were finally washed once with 0.1x SSC buffer. The resulting cDNAs released from chips were amplified with KAPA HiFi Hotstart Ready Mix (Roche, KK2602) with 0.8 μM cDNA-PCR primer. PCR reactions were conducted as: first incubation at 95 °C for 5 minutes, 15 cycles at 98 °C for 20 seconds, 58 °C for 20 seconds, 72 °C for 3 minutes and a final incubation at 72 °C for 5 minutes.

The concentrations of the resulting PCR products were quantified by Qubit™ dsDNA Assay Kit (Thermo, Q32854). A total of 20 ng of DNA were then fragmented with in-house Tn5 transposase at 55 °C for 10 minutes. Then, the reactions were stopped by the addition of 0.02% SDS buffer and gently mixing at 37 °C for 5 minutes. Fragmentation products were amplified as described below: 25 μL of fragmentation product, 1x KAPA HiFi Hotstart Ready Mix and 0.3 μM Stereo-Library-F primer, 0.3 μM Stereo-Library-R primer in a total volume of 100 μl with the addition of nuclease-free H2O. The reaction was then run as: 1 cycle of 95 °C for 5 minutes, 13 cycle of (98 °C 20 seconds, 58 °C 20 seconds and 72 °C 30 seconds) and 1 cycle 72 °C for 5 minutes. PCR products were purified using the Ampure XP Beads (Vazyme, N411-03) (0.6x and 0.2x) for DNB generation and finally sequenced (paired-end 50 bp) on a MGISEQ-2000RS sequencer.

### Raw Stereo-seq data processing

Fastq files were generated using a MGISEQ-2000RS sequencer, and the raw data was processed according to the Stereo-seq method(Chen A. et al., 2021). CID and MID are contained in the forward reads (CID: 1-25 bp, MID: 26-35 bp) while the reverse reads consist of the cDNA sequences. CID sequences on the forward reads were first mapped to the designed coordinates of the *in situ* captured chip, allowing 1 base mismatch to correct for sequencing and PCR errors. Reads with MID containing either N bases or more than 2 bases with quality scores lower than 10 were filtered out. CID and MID associated with each read were appended to each read header. Retained reads were then aligned to *Arabidopsis* genome (TAIR10) using STAR(Dobin et al., 2013) and mapped reads with MAPQ 10 were counted and annotated to their corresponding genes using an in-house script (available at https://github.com/BGIResearch/handleBam). MID with the same CID and the same gene locus were collapsed, allowing 1 mismatch to correct for sequencing and PCR errors. Finally, this information was used to generate a CID-containing expression profile matrix.

### Single cell data acquisition

To obtain single-cell level transcriptome data, we used the lasso tool provided in the visualization system of Stereomics to directly extract single-cell coordinate information and MID count matrix according to the bright field below MID signals. With every cell extracted, an information file was generated with bin1 size. We summed up the MID counts of the same gene with different X-Y coordinates in one information file and obtained the overall MID count for one single cell. After the generation of all single-cell information files, we processed the files to be suitable for subsequent analyses.

### Quality control of single-cell Stereo-seq data

Processing of the MID count matrix obtained from Stereo-seq data was implemented using the R package Seurat (version 4.0.0). To remove low-quality cells, we applied criteria to filter out cells with gene numbers no more than 200 or higher than 5000. Furthermore, we discarded low-quality cells with a high percentage of mitochondrial genes (>10%) to avoid perforated cells which lose cytoplasmic RNAs(Rich-Griffin et al., 2020). After filtering, the remaining 871 single cells with 14,425 genes were included in the downstream analyses.

### PCA analysis

The base R function prcomp is used to perform Principal Component Analysis (PCA). We obtained three-dimensional information of three principle components in our data set and visualized the spatial relationship between the different cell types of each leaf.

### Expression pattern analysis

We divided each leaf into seven parts, specifically, the main vein and the surrounding three leaf parts of equal length. The symmetrical leaf parts according to main vein were merged, which remains four unique areas in a leaf. The four parts of a single leaf were designated as Area 0, 1, 2 and 3, from main vein to leaf edge and were used for expression pattern analysis. The pattern analysis was performed utilizing the R package Mfuzz (2.50.0) with the mean expression of genes in four leaf areas.

### Trajectory analysis

Provided with spatial information of each single cell, the development direction of different cell types is certain, which is overall from Area 0 to 3. We used the R package Monocle (2.18.0)(Qiu et al., 2017) to carry out pseudotime analyses for vascular cell development and epidermal cell differentiation. For the exploration of all vasculature-related cells in our data, we selected DEGs between Area 0 and 3 cells that were detected in a minimum fraction of 0.14 and p-value < 0.05. The DEGs of epidermal cells and guard cells that were detected in a minimum fraction of 0.08 and p-value < 0.05 were chosen for the other analysis. Then, we reduced our data to two components using the method “DDRTree”. Cells were ordered along the developmental paths and visualized in two-dimensional space. Heatmaps were used to demonstrate the gene expression that differs in cells. The data used in plotting the heat maps was subsequently used for GO biological process analysis.

### GO enrichment analysis

GO enrichment analysis was performed using R package clusterProfiler(Yu et al., 2012) with TAIR10 annotation as the background. The smaller the *P*-value is, the more the GO term is significantly enriched.

## Supporting information

Supplemental Table 1, and related to Figure 1

## ACKNOWLEDGMENTS

This research was supported by Guangdong Provincial Key Laboratory of Genome Read and Write (No. 2017B030301011), and Shenzhen Key Laboratory of Single-Cell Omics (No. ZDSYS20190902093613831). We also sincerely thank the support provided by China National GeneBank (CNGB).

## AUTHOR CONTRIBUTIONS

X.X., Y.G., J.W., K.X., and H.-X.S. designed and supervised the study. K.X., H.-X.S., and Y.Z. designed the experiment. Y.Z., G.L., Z.C., and R.Y. performed the library preparation and sequencing. J.L., R.C., S.H. and H.-X.S. performed bioinformatics analysis. J.X. provided technical support. J.W., Q.X., Y.L., A.C., L.L., Y.Y., and H.Y. gave the relevant advice. K.X., H.-X.S., J.-M.L. J.L., G.L., Z.C. and R.C. wrote the manuscript. K.X., H.-X.S., X. X. and Y. G. participated in the manuscript editing and discussion. All authors edited and approved the manuscript.

## DATA AVAILABILITY

The high-throughput sequencing data that support the findings of this study have been deposited into CNGB Sequence Archive(Guo et al., 2020) of CNGBdb(Chen et al., 2020) with accession number CNP0001923.

## Supplementary Information

**Figure S1.**
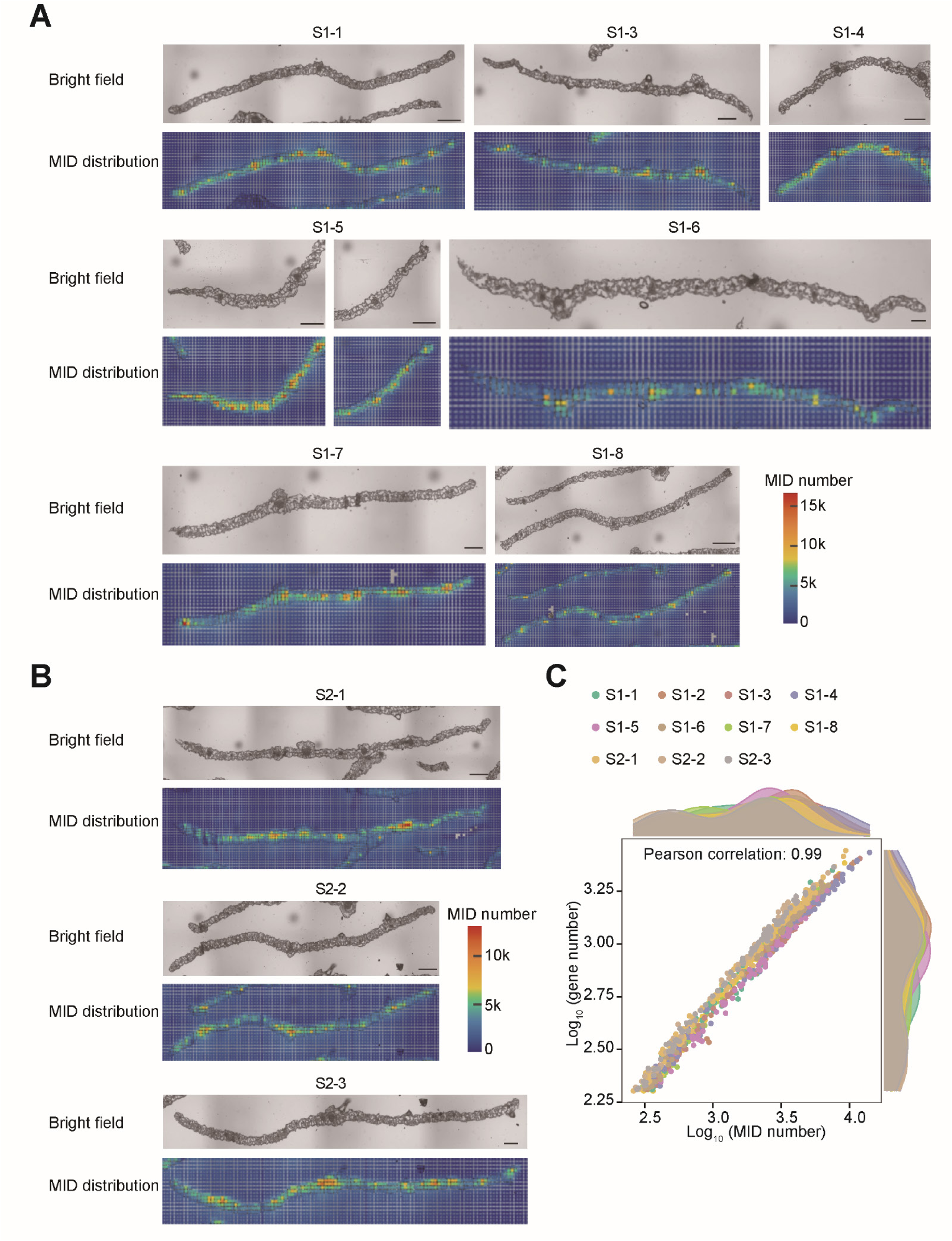
MID distribution in 11 *Arabidopsis thaliana* cauline leaves, Related to Figure 1. (A and B) Bright field image and MID distribution in *Arabidopsis* cauline leaves from Section 1 (S1, A) and Section 2 (S2, B). The scale bar is 500 μm. The color bar represents the number of MIDs. (C) Scatterplot showing gene number and MID number per cell in 11 leaves. All points are summarized and shown as density plots along each axis.

**Figure S2.**
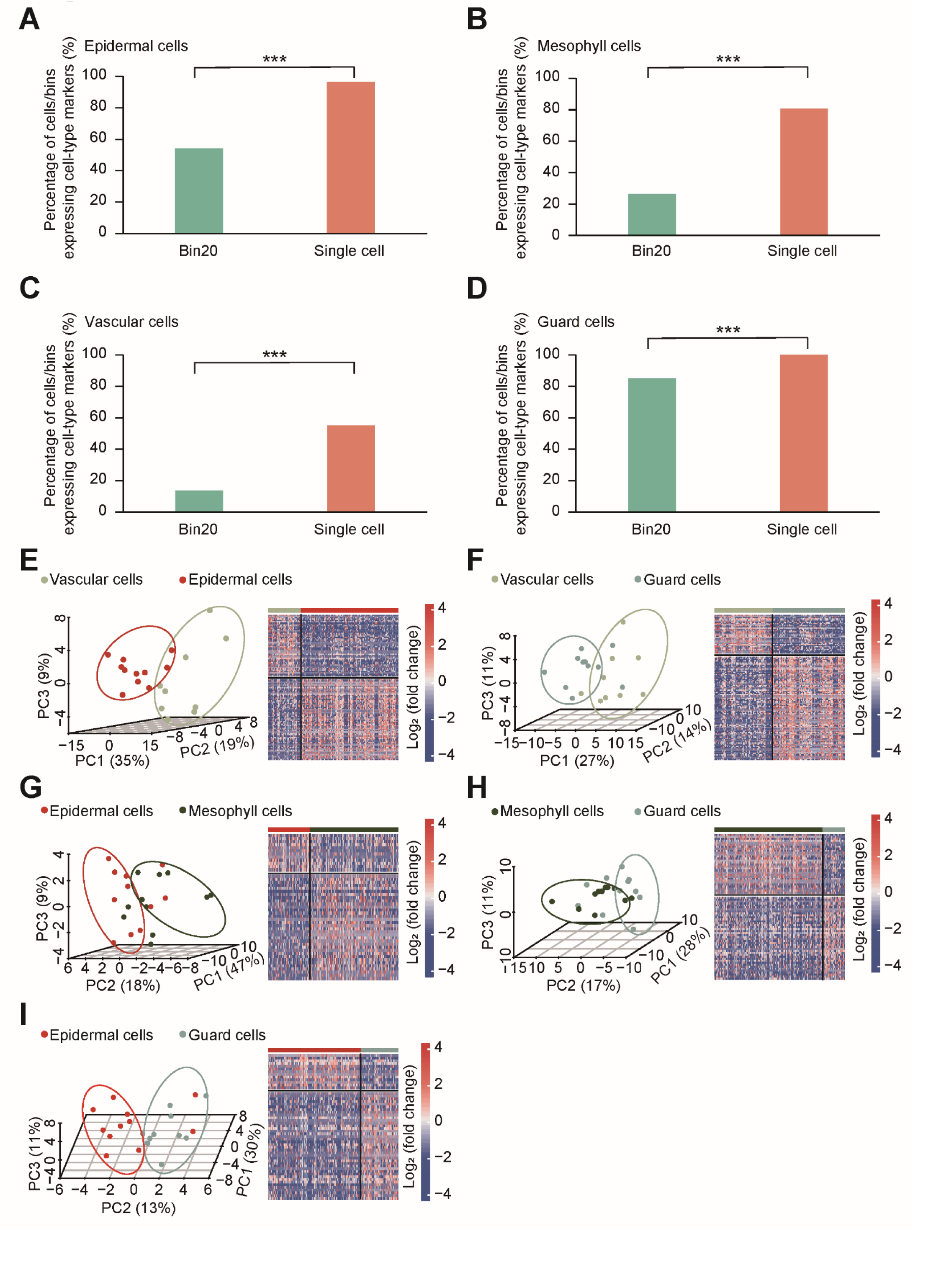
Comparison of transcriptome profiles between bins and single cells and technical reproducibility of cell types from different cauline leaves, Related to Figure 2. (A-D) Percentage of cells/bins expressing markers of epidermal cells (A), mesophyll cells (B), vascular cells (C) and guard cells (D). (E-I) PCA plot of vascular cells and epidermal cells (E), vascular cells and guard cells (F), epidermal cells and mesophyll cells (G), mesophyll cells and guard cells (H), and epidermal cells and guard cells (I) in different leaves (left). The heatmaps showing DEGs between the above cell types are shown on the right. Asterisks indicate statistically significant differences: (***) P < 0.001.

**Figure S3.**
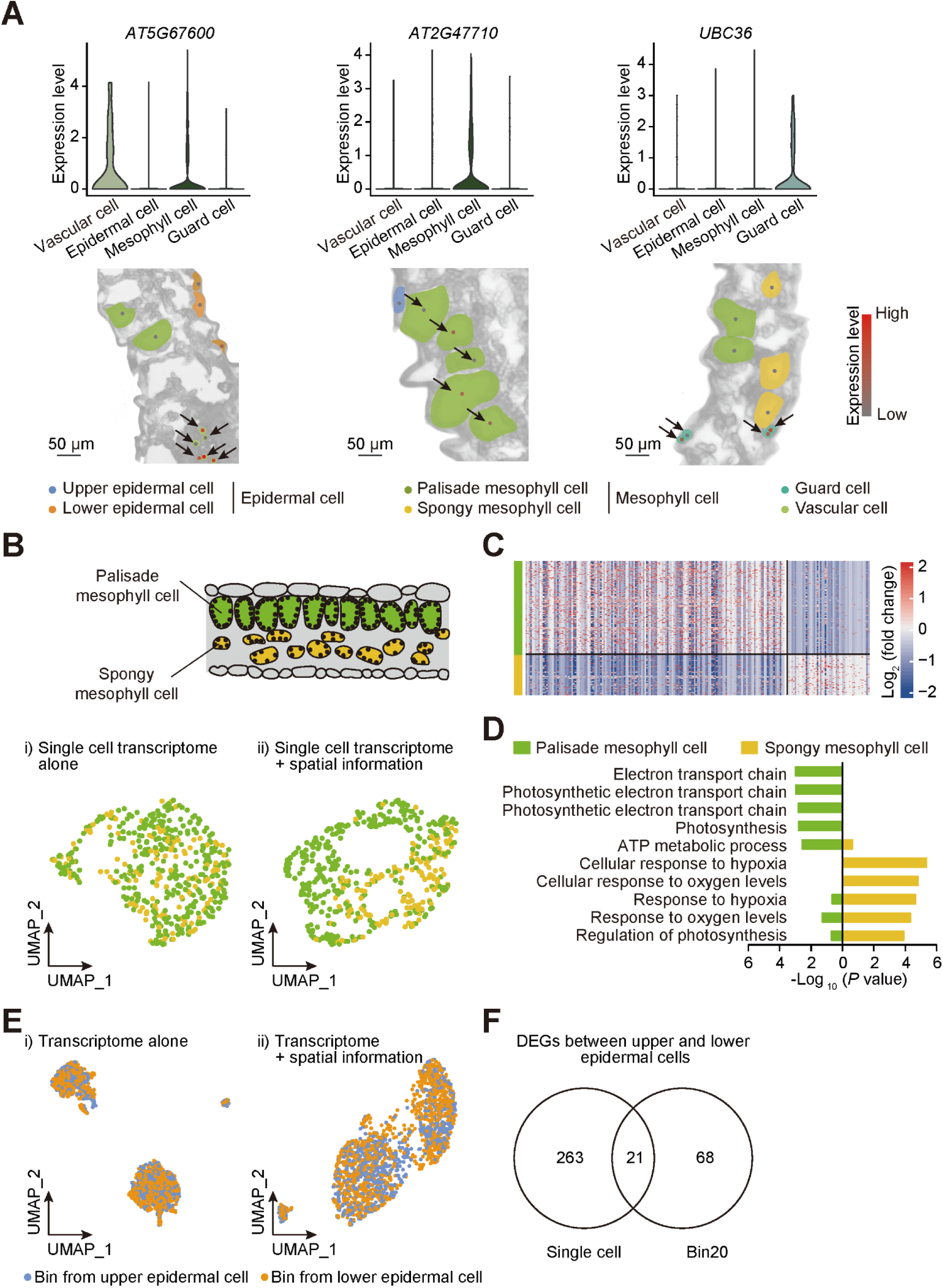
Identification and characterization of cell type and cell sub-type, Related to Figure 3. (A) Validation of cell types using cell-type specific marker genes (top panel). Representative images showing expression levels of marker genes in the corresponding cells in the bright field (lower panel). (B) Spatial information allows separation between palisade and spongy mesophyll cells. Distribution of palisade and spongy mesophyll cells: i) without spatial information (using variable genes of all mesophyll cells); and ii) with spatial information (using DEGs between palisade and spongy mesophyll cells). Green dots represent palisade mesophyll cell and yellow dots represent spongy mesophyll cell. (C) DEGs between palisade mesophyll cell and spongy mesophyll cell. Blue and red represent log2-transformed fold change < 0 and > 0, respectively. (D) GO enrichment analysis of DEGs between palisade mesophyll cell and spongy mesophyll cell. (E) UMAP projection plot of bins from epidermal cells using: i) variable genes of bins from epidermal cells; and ii) differentially expressed genes (DEGs) between bins from upper and lower epidermal cells. (F) Only 21 DEGs were overlapped between upper and lower epidermal cells using single-cell method and Bin20 method.

**Figure S4.**
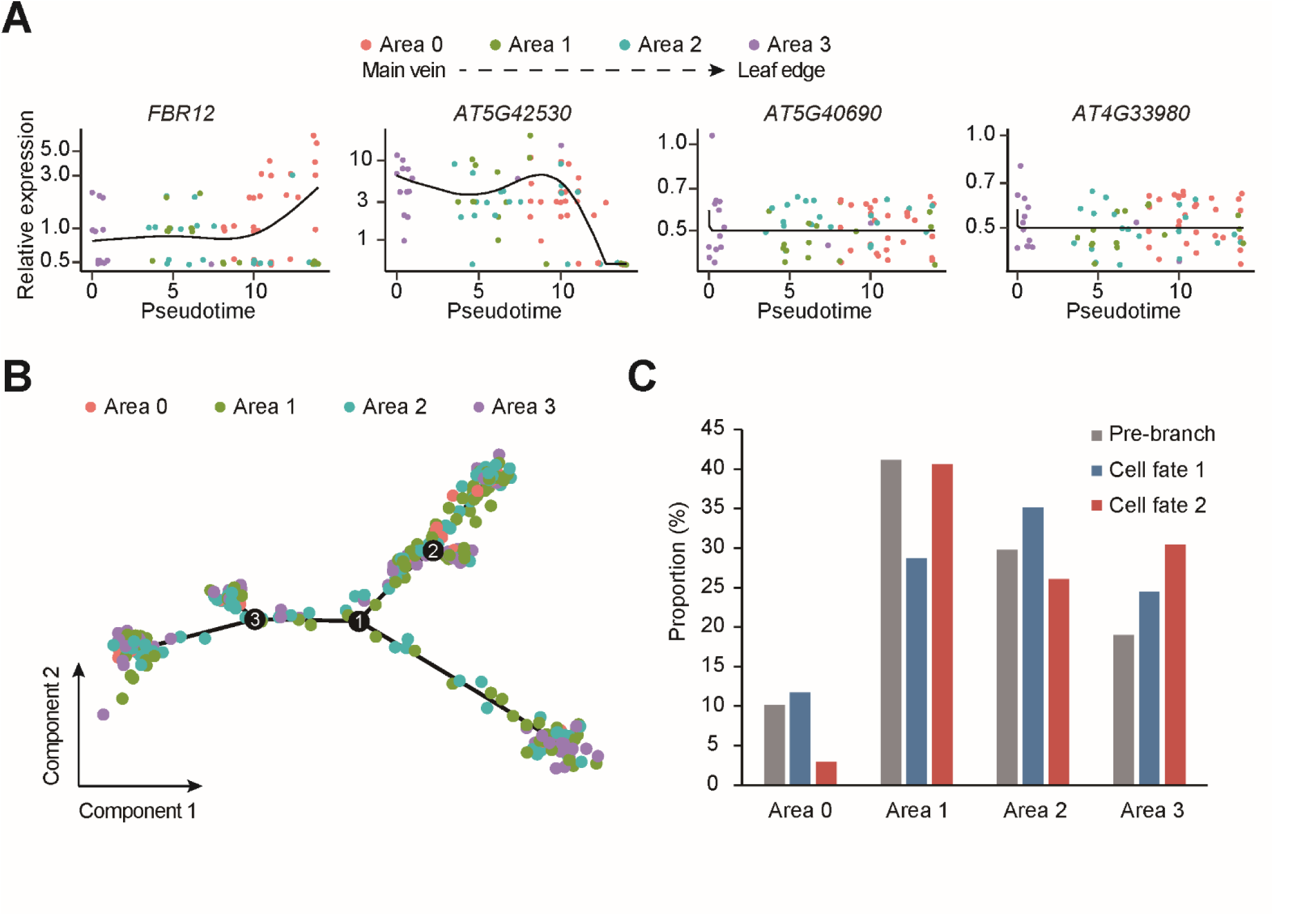
Spatially resolved expression patterns of vascular cells and spatial distribution of guard cells and epidermal cells, Related to Figure 5. (A) Gene expression kinetics along pseudotime progression for representative genes from vascular cells. (B) Distribution of guard cells and epidermal cells from each area on the pseudotime trajectory. (C) Statistics of spatial distribution of guard cells and epidermal cells in each area from each branch.

**Table S1 Statistics of single cells detected by scStereo-seq from 11 cauline leaves, Related to Figure 1**

## REFERENCES

Bezrutczyk, M., Zollner, N.R., Kruse, C.P.S., Hartwig, T., Lautwein, T., Kohrer, K., Frommer, W.B., and Kim, J.Y. (2021). Evidence for phloem loading via the abaxial bundle sheath cells in maize leaves. Plant Cell 33, 531–547.

Brown, N.J., Palmer, B.G., Stanley, S., Hajaji, H., Janacek, S.H., Astley, H.M., Parsley, K., Kajala, K., Quick, W.P., Trenkamp, S., et al. (2010). C acid decarboxylases required for C photosynthesis are active in the mid-vein of the C species Arabidopsis thaliana, and are important in sugar and amino acid metabolism. Plant J 61, 122–133.

Chen A., Chen A., Chen A., Chen A., Chen A., Chen A., and Chen A. (2021). Large field of view-spatially resolved transcriptomics at nanoscale resolution. bioRxiv.

Chen, F.Z., You, L.J., Yang, F., Wang, L.N., Guo, X.Q., Gao, F., Hua, C., Tan, C., Fang, L., Shan, R.Q., et al. (2020). CNGBdb: China National GeneBank DataBase. Yi Chuan 42, 799–809.

Chen, K.H., Boettiger, A.N., Moffitt, J.R., Wang, S., and Zhuang, X. (2015). RNA imaging. Spatially resolved, highly multiplexed RNA profiling in single cells. Science 348, aaa6090.

Cho, C.S., Xi, J., Si, Y., Park, S.R., Hsu, J.E., Kim, M., Jun, G., Kang, H.M., and Lee, J.H. (2021). Microscopic examination of spatial transcriptome using Seq-Scope. Cell 184, 3559–3572 e3522.

Dobin, A., Davis, C.A., Schlesinger, F., Drenkow, J., Zaleski, C., Jha, S., Batut, P., Chaisson, M., and Gingeras, T.R. (2013). STAR: ultrafast universal RNA-seq aligner. Bioinformatics (Oxford, England) 29, 15–21.

Eng, C.L., Lawson, M., Zhu, Q., Dries, R., Koulena, N., Takei, Y., Yun, J., Cronin, C., Karp, C., Yuan, G.C., et al. (2019). Transcriptome-scale super-resolved imaging in tissues by RNA seqFISH. Nature 568, 235–239.

Frenette Charron, J.B., Breton, G., Badawi, M., and Sarhan, F. (2002). Molecular and structural analyses of a novel temperature stress-induced lipocalin from wheat and Arabidopsis. FEBS Lett 517, 129–132.

Gao, Z., Shen, W., and Chen, G. (2018). Uncovering C4-like photosynthesis in C3 vascular cells. J Exp Bot 69, 3531–3540.

Geng, P., Zhang, S., Liu, J., Zhao, C., and Zhao, Q. (2019). MYB20, MYB42, MYB43 and MYB85 Regulate Phenylalanine and Lignin Biosynthesis during Secondary Cell Wall Formation. Plant Physiology 182, pp.01070.02019.

Giacomello, S., Salmen, F., Terebieniec, B.K., Vickovic, S., Navarro, J.F., Alexeyenko, A., Reimegard, J., McKee, L.S., Mannapperuma, C., Bulone, V., et al. (2017). Spatially resolved transcriptome profiling in model plant species. Nat Plants 3, 17061.

Glover, B.J., Airoldi, C.A., and Moyroud, E. (2016). Epidermis: Outer Cell Layer of the Plant. eLS.

Guo, X., Chen, F., Gao, F., Li, L., Liu, K., You, L., Hua, C., Yang, F., Liu, W., Peng, C., et al. (2020). CNSA: a data repository for archiving omics data. Database (Oxford) 2020.

Gurazada, S.G.R., Cox, K.L., Czymmek, K.J., and Meyers, B.C. (2021). Space: the final frontier - achieving single-cell, spatially resolved transcriptomics in plants. Emerg Top Life Sci.

Hibberd, J.M., and Quick, W.P. (2002). Characteristics of C4 photosynthesis in stems and petioles of C3 flowering plants. Nature 415, 451–454.

Ischebeck, T., Werner, S., Krishnamoorthy, P., Lerche, J., Meijon, M., Stenzel, I., Lofke, C., Wiessner, T., Im, Y.J., Perera, I.Y., et al. (2013). Phosphatidylinositol 4,5-bisphosphate influences PIN polarization by controlling clathrin-mediated membrane trafficking in Arabidopsis. The Plant cell 25, 4894–4911.

Ishihara, S., Takabayashi, A., Ido, K., Endo, T., Ifuku, K., and Sato, F. (2007). Distinct functions for the two PsbP-like proteins PPL1 and PPL2 in the chloroplast thylakoid lumen of Arabidopsis. Plant Physiol 145, 668–679.

Jiao, Y., Tausta, S.L., Gandotra, N., Sun, N., Liu, T., Clay, N.K., Ceserani, T., Chen, M., Ma, L., Holford, M., et al. (2009). A transcriptome atlas of rice cell types uncovers cellular, functional and developmental hierarchies. Nat Genet 41, 258–263.

Kawamura, Y., and Uemura, M. (2003). Mass spectrometric approach for identifying putative plasma membrane proteins of Arabidopsis leaves associated with cold acclimation. Plant J 36, 141–154.

Keren, N., Ohkawa, H., Welsh, E.A., Liberton, M., and Pakrasi, H.B. (2005). Psb29, a conserved 22-kD protein, functions in the biogenesis of Photosystem II complexes in Synechocystis and Arabidopsis. Plant Cell 17, 2768–2781.

Kim, J.-Y., Kim, J.-Y., Kim, J.-Y., Kim, J.-Y., Kim, J.-Y., Kim, J.-Y., Kim, J.-Y., and Kim, J.-Y. (2021). Distinct identities of leaf phloem cells revealed by single cell transcriptomics. Plant Cell.

Kimura, M., Yamamoto, Y.Y., Seki, M., Sakurai, T., Sato, M., Abe, T., Yoshida, S., Manabe, K., Shinozaki, K., and Matsui, M. (2003). Identification of Arabidopsis genes regulated by high light-stress using cDNA microarray. Photochem Photobiol 77, 226–233.

Kubo, M., Nishiyama, T., Tamada, Y., Sano, R., Ishikawa, M., Murata, T., Imai, A., Lang, D., Demura, T., Reski, R., et al. (2019). Single-cell transcriptome analysis of Physcomitrella leaf cells during reprogramming using microcapillary manipulation. Nucleic Acids Res 47, 4539–4553.

Lee, J.H., Daugharthy, E.R., Scheiman, J., Kalhor, R., Yang, J.L., Ferrante, T.C., Terry, R., Jeanty, S.S., Li, C., Amamoto, R., et al. (2014). Highly multiplexed subcellular RNA sequencing in situ. Science 343, 1360–1363.

Liu, Y., Yang, M., Deng, Y., Su, G., Enninful, A., Guo, C.C., Tebaldi, T., Zhang, D., Kim, D., Bai, Z., et al. (2020a). High-Spatial-Resolution Multi-Omics Sequencing via Deterministic Barcoding in Tissue. Cell 183, 1665–1681 e1618.

Liu, Z., Zhou, Y., Guo, J., Li, J., Tian, Z., Zhu, Z., Wang, J., Wu, R., Zhang, B., Hu, Y., et al. (2020b). Global Dynamic Molecular Profiling of Stomatal Lineage Cell Development by Single-Cell RNA Sequencing. Mol Plant 13, 1178–1193.

Lopez-Anido, C.B., Vaten, A., Smoot, N.K., Sharma, N., Guo, V., Gong, Y., Anleu Gil, M.X., Weimer, A.K., and Bergmann, D.C. (2021). Single-cell resolution of lineage trajectories in the Arabidopsis stomatal lineage and developing leaf. Dev Cell 56, 1043–1055 e1044.

Martinez, C.C., Li, S., Woodhouse, M.R., Sugimoto, K., and Sinha, N.R. (2021). Spatial transcriptional signatures define margin morphogenesis along the proximal-distal and medio-lateral axes in tomato (Solanum lycopersicum) leaves. Plant Cell 33, 44–65.

Maugarny-Cales, A., and Laufs, P. (2018). Getting leaves into shape: a molecular, cellular, environmental and evolutionary view. Development 145.

Miki, Y., Takahashi, D., Kawamura, Y., and Uemura, M. (2019). Temporal proteomics of Arabidopsis plasma membrane during cold- and de-acclimation. J Proteomics 197, 71–81.

Morales, R., Charon, M.H., Kachalova, G., Serre, L., Medina, M., Gomez-Moreno, C., and Frey, M. (2000). A redox-dependent interaction between two electron-transfer partners involved in photosynthesis. EMBO Rep 1, 271–276.

Nichterwitz, S., Chen, G., Aguila Benitez, J., Yilmaz, M., Storvall, H., Cao, M., Sandberg, R., Deng, Q., and Hedlund, E. (2016). Laser capture microscopy coupled with Smart-seq2 for precise spatial transcriptomic profiling. Nat Commun 7, 12139.

Peng, L., Fukao, Y., Fujiwara, M., Takami, T., and Shikanai, T. (2009). Efficient operation of NAD(P)H dehydrogenase requires supercomplex formation with photosystem I via minor LHCI in Arabidopsis. Plant Cell 21, 3623–3640.

Qiu, X., Mao, Q., Tang, Y., Wang, L., Chawla, R., Pliner, H.A., and Trapnell, C. (2017). Reversed graph embedding resolves complex single-cell trajectories. Nat Methods 14, 979–982.

Rhee, S.Y., Birnbaum, K.D., and Ehrhardt, D.W. (2019). Towards Building a Plant Cell Atlas. Trends Plant Sci 24, 303–310.

Rich-Griffin, C., Stechemesser, A., Finch, J., Lucas, E., Ott, S., and Schafer, P. (2020). Single-Cell Transcriptomics: A High-Resolution Avenue for Plant Functional Genomics. Trends Plant Sci 25, 186–197.

Rodriques, S.G., Stickels, R.R., Goeva, A., Martin, C.A., Murray, E., Vanderburg, C.R., Welch, J., Chen, L.M., Chen, F., and Macosko, E.Z. (2019). Slide-seq: A scalable technology for measuring genome-wide expression at high spatial resolution. Science 363, 1463–1467.

Ruan, Y.L. (2014). Sucrose metabolism: gateway to diverse carbon use and sugar signaling. Annu Rev Plant Biol 65, 33–67.

Shaw, R., Tian, X., and Xu, J. (2021). Single-Cell Transcriptome Analysis in Plants: Advances and Challenges. Mol Plant 14, 115–126.

Srivatsan, S.R., Regier, M.C., Barkan, E., Franks, J.M., Packer, J.S., Grosjean, P., Duran, M., Saxton, S., Ladd, J.J., Spielmann, M., et al. (2021). Embryo-scale, single-cell spatial transcriptomics. Science 373, 111–117.

Stahl, P.L., Salmen, F., Vickovic, S., Lundmark, A., Navarro, J.F., Magnusson, J., Giacomello, S., Asp, M., Westholm, J.O., Huss, M., et al. (2016). Visualization and analysis of gene expression in tissue sections by spatial transcriptomics. Science 353, 78–82.

Stickels, R.R., Murray, E., Kumar, P., Li, J., Marshall, J.L., Di Bella, D.J., Arlotta, P., Macosko, E.Z., and Chen, F. (2021). Highly sensitive spatial transcriptomics at near-cellular resolution with Slide-seqV2. Nat Biotechnol 39, 313–319.

Svozil, J., Gruissem, W., and Baerenfaller, K. (2015). Proteasome targeting of proteins in Arabidopsis leaf mesophyll, epidermal and vascular tissues. Front Plant Sci 6, 376.

Tian, C., Du, Q., Xu, M., Du, F., and Jiao, Y. (2020). Single-nucleus RNA-seq resolves spatiotemporal developmental trajectories in the tomato shoot apex. bioRxiv.

Tian, C., Wang, Y., Yu, H., He, J., Wang, J., Shi, B., Du, Q., Provart, N.J., Meyerowitz, E.M., and Jiao, Y. (2019). A gene expression map of shoot domains reveals regulatory mechanisms. Nat Commun 10, 141.

Tsukaya, H. (2013). Leaf development. Arabidopsis Book 11, e0163.

Vickovic, S., Eraslan, G., Salmen, F., Klughammer, J., Stenbeck, L., Schapiro, D., Aijo, T., Bonneau, R., Bergenstrahle, L., Navarro, J.F., et al. (2019). High-definition spatial transcriptomics for in situ tissue profiling. Nat Methods 16, 987–990.

Wang, J., Sun, N., Zhang, F., Yu, R., Chen, H., Deng, X.W., and Wei, N. (2020). SAUR17 and SAUR50 Differentially Regulate PP2C-D1 during Apical Hook Development and Cotyledon Opening in Arabidopsis. Plant Cell 32, 3792–3811.

Wang, X., Allen, W.E., Wright, M.A., Sylwestrak, E.L., Samusik, N., Vesuna, S., Evans, K., Liu, C., Ramakrishnan, C., Liu, J., et al. (2018). Three-dimensional intact-tissue sequencing of single-cell transcriptional states. Science 361.

Xia, C., Fan, J., Emanuel, G., Hao, J., and Zhuang, X. (2019). Spatial transcriptome profiling by MERFISH reveals subcellular RNA compartmentalization and cell cycle-dependent gene expression. Proc Natl Acad Sci U S A 116, 19490–19499.

Yabuta, S., Ifuku, K., Takabayashi, A., Ishihara, S., Ido, K., Ishikawa, N., Endo, T., and Sato, F. (2010). Three PsbQ-like proteins are required for the function of the chloroplast NAD(P)H dehydrogenase complex in Arabidopsis. Plant Cell Physiol 51, 866–876.

Yu, G., Wang, L.G., Han, Y., and He, Q.Y. (2012). clusterProfiler: an R package for comparing biological themes among gene clusters. OMICS 16, 284–287.

Zhang, T.Q., Chen, Y., and Wang, J.W. (2021). A single-cell analysis of the Arabidopsis vegetative shoot apex. Dev Cell 56, 1056–1074 e1058.

